# Wind history shapes olfactory search response in free flying *Drosophila melanogaster*

**DOI:** 10.64898/2026.04.05.716000

**Authors:** Jaleesa Houle, Austin Lopez, Floris van Breugel

**Author notes:** Corresponding author: Floris van Breuge.

## Abstract

The ability of flying insects to locate distant food and mates by tracking odor plumes through turbulent and unsteady flow represents a remarkable feat of sensorimotor integration. Successful navigation requires not only extracting a reliable directional estimate from an intermittent olfactory signal, but also contending with the challenging dynamics of variable winds. While prior work has established that insects integrate the history of odor encounters to shape search decisions, whether they also retain a working memory of recently experienced wind conditions has remained unknown. Here, we use optogenetics combined with controlled wind perturbations in a free-flight wind tunnel to investigate how wind history modulates the olfactory search behavior of *Drosophila melanogaster*. By introducing lateral “gust” flow via auxiliary fans and independently delivering olfactory stimuli, we show that the wind experienced *during* an olfactory stimulus shapes both the immediate surge response and the subsequent spatial search. Flies that received an olfactory stimulus while being displaced by a crosswind gust were significantly more likely to return to the gust zone during the post-stimulus search phase compared to flies that received the same odor cue in steady laminar flow. Meanwhile, surge responses and course directions exhibited during search indicate that moment-to-moment flight kinematics may be driven more by instantaneous flow. These results reveal that wind experience is tracked in addition to olfactory experience, and provide evidence that *Drosophila* maintain a short-term working memory of ambient wind conditions to guide olfactory navigation.

## INTRODUCTION

Locating distant sources of food and mates through olfactory cues is among the most ecologically consequential behaviors performed by flying insects. Whether it is a male moth traveling kilometers to find a pheromone-releasing female or a mosquito seeking a host, these olfactory-driven behaviors shape ecosystems and influence the spread of vector-borne diseases (Murlis et al., 1992; Cardé and Willis, 2008; Dekker and Cardé, 2011; Van Breugel et al., 2015). In nearly all such contexts, odor cues are not transmitted as smooth, continuous gradients but are fragmented into intermittent, stochastic filaments that an animal must detect, integrate, and act upon in real time (Murlis et al., 1992; Celani et al., 2014). Understanding how insects solve this navigational problem has broad implications for the development of autonomous chemical-sensing platforms, as well as for ecological and public health efforts including trap optimization and disease vector control (Bau and Cardé, 2015; Carey and Carlson, 2011; Anderson et al., 2020).

A well-established behavioral motif for insect odor plume tracking, developed through decades of work with moths, flies, and other taxa, describes an iterative two-phase strategy known as cast-and-surge (Kennedy and Marsh, 1974; Kennedy, 1983; Van Breugel and Dickinson, 2014). Upon encountering a filament of attractive odor, a flying insect executes a rapid upwind surge. After losing the plume, the animal transitions into crosswind casting, sweeping back and forth to reacquire the odor signal (Cardé and Willis, 2008; Mafra-Neto and Cardé, 1994; Vickers and Baker, 1994; Van Breugel and Dickinson, 2014). Flies accomplish this behavior through multi-modal sensory integration, including optic flow via visual feedback (Chow et al., 2011; Chow and Frye, 2008) and bilateral comparison of odor concentration across the antennae to estimate source direction (Duistermars et al., 2009; Murthy, 2024; Kadakia et al., 2022). Although visual cues and olfactory stimuli are both important for executing successful search bouts, the ambient wind also plays an important role as it advects the odor from its source. Wind direction itself is encoded centrally in the fly brain through bilateral comparison of antennal mechanosensory signals and decoded by dedicated projection neurons in the fly brain (Suver et al., 2019). Walking insects employ a related strategy, combining odor-gated upwind running with local area search triggered by odor offset, similarly relying on antennal mechanoreception for wind orientation (Álvarez Salvado et al., 2018; Willis et al., 2011).

The effectiveness of the cast-and-surge strategy depends critically on wind conditions, and natural wind environments are far more complex than the laminar flows of standard laboratory wind tunnels. In laminar flow, odor plumes form predictable ribbon-like structures that insects exploit efficiently using stereotyped reflexes (van Breugel and Dickinson, 2014; Cardé and Willis, 2008). As turbulence increases and wind direction becomes variable, plumes disperse and meander, fundamentally altering the statistics of odor encounters and subsequently reshaping navigational behavior (Talley et al., 2023; Murlis et al., 1992). Near-surface wind direction variability in natural environments can fluctuate substantially on timescales of 1–10 minutes, suggesting that real-world plume tracking routinely occurs under conditions poorly approximated by conventional wind tunnel paradigms (Houle and van Breugel, 2023, 2026). A more recent finding in *Drosophila* has shown that flies rapidly assess ambient wind within approximately 100 ms of odor onset and select qualitatively distinct search strategies accordingly: cast-and-surge in laminar wind, and a stereotyped “sink and circle” local search, characterized by rhythmic unidirectional turns in still air (Stupski and van Breugel, 2024). This finding demonstrates that wind information does not merely direct upwind flight, but actively gates the downstream navigational state.

While instantaneous conditions clearly shape insect search strategies, accumulating evidence indicates that insects also integrate sensory information across time, drawing on the history of prior encounters to guide navigation. Pang et al. (2018b) demonstrated that the strength of odor-triggered upwind turns in flying *Drosophila* and *Aedes aegypti* is modulated by the history of prior odor encounters within a trajectory, with later encounters in a sequence eliciting progressively weaker upwind turns, consistent with short-term memory for odor experience accumulating over multi-encounter timescales. In walking *Drosophila*, the timing of prior odor encounters similarly biases the direction of stochastic saccades upwind, with encounter frequency additionally modulating walk–stop transition rates (Álvarez Salvado et al., 2018). Deep reinforcement learning models trained to navigate turbulent plumes corroborate the importance of sensory memory, showing that longer integration timescales improve tracking performance, particularly when wind direction shifts (Singh et al., 2023).

While memory for odor experience has been experimentally demonstrated, whether insects also retain and use a working memory of recently experienced wind conditions to shape subsequent search decisions has not been established. This knowledge gap arises in part from the difficulty of decoupling odor and wind stimuli in conventional experiments. Optogenetics offers a way to overcome this barrier by enabling direct, spatiotemporally precise control of olfactory input, independent of flow (Klapoetke et al., 2014; Stupski and van Breugel, 2024). In this study, we exploit optogenetic stimulation alongside controlled flow perturbations—lateral “gusts” produced by auxiliary fans placed downwind within our wind tunnel— to recreate the kind of unsteady wind environment that flies might encounter during a natural outdoor plume-tracking bout. By combining high-resolution 3-D flight tracking with independently controlled wind and odor stimulation, we compare the trajectories of flies that experienced distinct flow histories at the moment of olfactory stimulus. Our findings reveal that the wind experienced during an olfactory stimulus influences where a fly subsequently searches for the odor source, while course direction preferences may be dictated more by instantaneous flow. Taken together, this work provides evidence that *Drosophila* retain a short-term memory of recent wind conditions during olfactory navigation.

## MATERIALS AND METHODS

### Wind Tunnel Illumination and Trajectory Tracking

We used the same wind tunnel illumination, cameras, and trajectory tracking protocol described previously in Stupski and van Breugel (2024). In brief, 12 Basler cameras (acA720, Basler AG, Ahrensburg, Germany) operating at 100 fps are stationed above the tunnel (Figure 1A). The cameras are equipped with IR-pass filters to track trajectories using infrared light. The 3-D trajectories are tracked in real-time and recorded using open source software (Braid (Straw et al., 2011)). The wind tunnel walls and floor are illuminated with arrays of infrared and blue LEDs to light the interior, while the roof is fitted with white LEDs to provide orange-wavelength light for photoreceptor re-isomerization. Walls other than the roof are lined with diffusing film for uniform lighting, and total illumination from the combined blue and white LEDs measured 425 lux.

**Figure 1:**
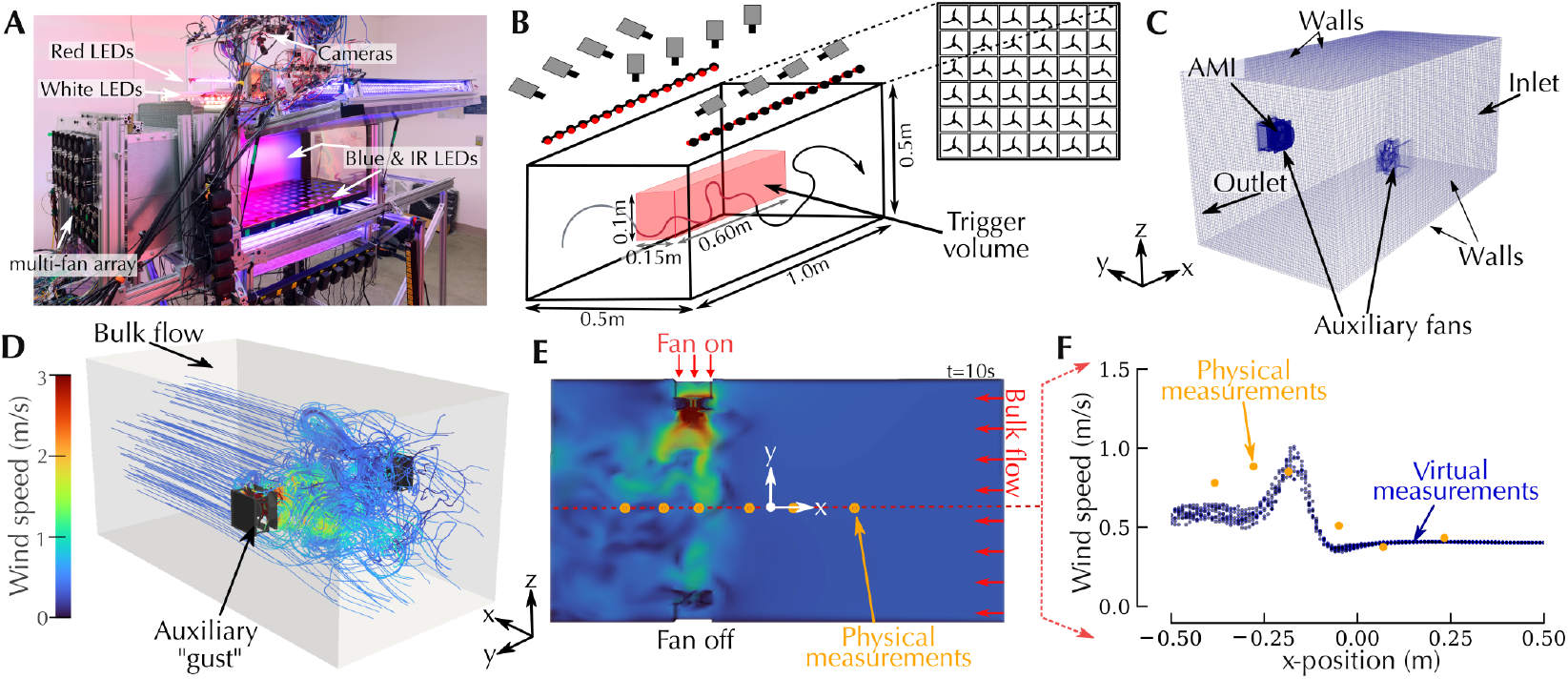
Physical measurements and CFD simulations demonstrate distinct flow regions within the wind tunnel. (A) Wind tunnel equipped with red, white, blue, and IR lights for controlling optogenetic responses in *Drosophila melanogaster*. (B) Schematic showing cameras, red lights, upwind fan array, and experiment trigger volume. When a fly enters the trigger volume, it receives either a 0.675s red light flash or no olfactory stimulus (sham). (C) Mesh and boundary conditions used in CFD simulations. Two auxiliary fans were modeled, with only one simulated as “on” to match the wind tunnel settings. The AMI (arbitrary mesh interface) is shown for the fan that was “on”, modeled as a moving mesh (see Methods for more details). (D) Example streamlines within the computational domain at *t* = 10s, colored by wind speed. (E) Instantaneous wind speed profile at *t* = 10s for altitudes 0.21 ≤ *z* ≤ 0.29m, corresponding to the auxiliary fan locations on the side walls. Color map is the same as in (D). (F) Comparison of mean wind speed for points taken along the *y*-centerline (red dotted line in E) between CFD virtual measurements (blue, 5 ≤ *t* ≤ 20s) and physical measurements (orange, *∼*2 minutes per location).

### Optogenetic Stimulus

The experimental paradigms for these experiments mirror those described previously (Stupski and van Breugel, 2024). Custom software (C++ and Python) was implemented via the Robot Operating System (ROS) to coordinate the timing of red light stimulus with trajectories moving through a pre-designated 3-D trigger zone within the tunnel (Figure 1B). For these experiments, flies that entered the trigger zone were randomly assigned a 675 ms pulse of light, referred to as a “flash” event, or no light, deemed “sham” events. The red light stimuli for these experiments were generated by arrays of Triple Bright RGB LEDs (SparkFun Electronics, Niwot, CO, USA), which line the top of both sides of the wind tunnel (Figure 1A-B). A 5-second refractory period was programmed after trigger events to prevent rapid reactivation. The red light intensity in the middle of the tunnel was 42 µW/m^2^ of 625 nm light, see (Stupski and van Breugel, 2024) for a more detailed characterization.

### Wind Tunnel Flow Measurements

The flow of air into the wind tunnel is driven by a multi-fan array (MFA) that consists of a 6 × 6 grid of individually controllable 80 mm computer fans (Lopez, 2024). Full details regarding our wind tunnel design and flow capabilities are discussed in Lopez (2024). For this experiment, we set the fan array to produce a uniform laminar flow (*U* = 0.4*m/s*). To create a “gust” scenario, we added auxiliary fans (Delta Electronics, AUB0812L-9X41, FAN AXIAL 80X25.4MM 12VDC) on the left and right side walls (Figure 1C) approximately 0.3 meters upwind of the outlet. The auxiliary fans were programmed to consecutively cycle on and off at 20 minute intervals, such that only one fan was active at a time. Between each cycle, we included a 20 minute interval in which both fans remained off.

We measured flow properties inside the wind tunnel with a 2-D ultrasonic anemometer (LI-550 TriSonica Mini), sampling at 10Hz after calibration in a high-damping enclosure. A custom aluminum frame enabled precise positioning and allowed the sensor to be oriented in the (x,y) plane along the tunnel length. Using this setup, wind speed and direction data were collected at six streamwise locations in the test chamber.

### Computational Fluid Dynamics (CFD) Simulations

To visualize flow characteristics within the wind tunnel with added auxiliary fans, we performed numerical simulations using the 3-D incompressible Navier-Stokes equations:

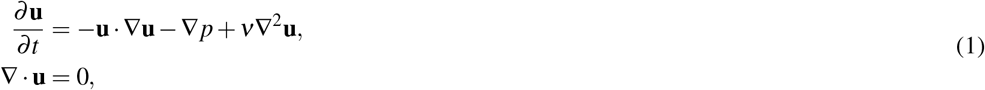

where **u** = **u**(**x**, *t*) is the velocity field over spatial reference frame **x** and time *t, p* is the pressure divided by density, and ν = 1 10^−5^ m^2^*/*s is the kinematic viscosity. The numerical solution was computed using rhoPimpleFOAM, a transient solver for incompressible flow of Newtonian fluids, available in the open-source CFD software OpenFOAM (Greenshields et al., 2018). The Reynolds-Averaged Navier-Stokes (RANS) approach with *k*-ω SST turbulence closure was employed to model turbulent fluctuations.

The computational domain, designed to represent the experimental wind tunnel section, measured 1.0 m×0.5 m×0.5m (length×width×height) as shown schematically in Figure 1B. A rotating fan was positioned within the domain to model the auxiliary “gust”, with a second stationary fan modeled opposite the “gust” (Figure 1C). The boundary conditions are summarized in Table 1.

**Table 1:**
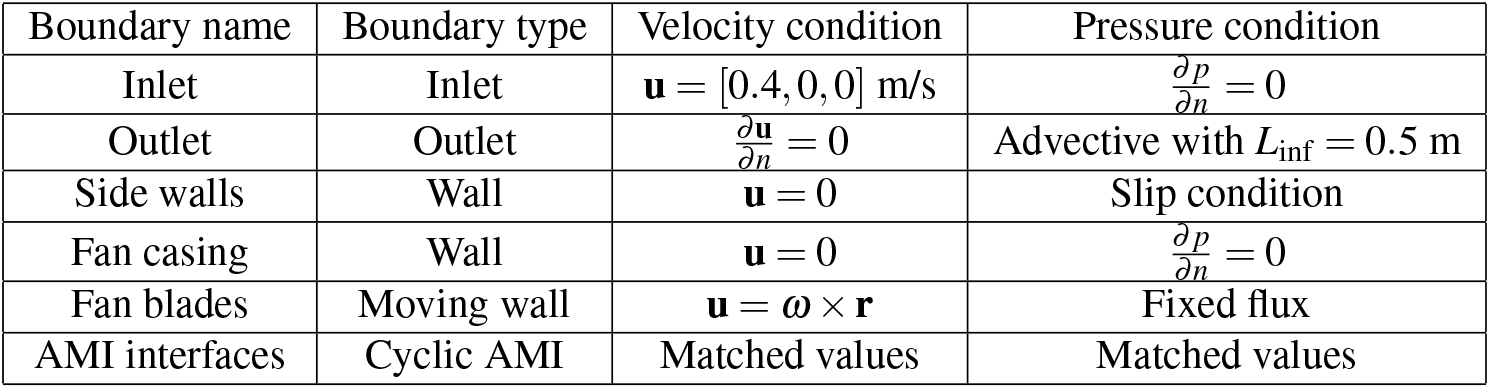
Boundary conditions applied in the wind tunnel simulation with auxiliary fan.

| Boundary name | Boundary type | Velocity condition | Pressure condition |
| --- | --- | --- | --- |
| Inlet | Inlet | $\mathbf{u} = [0.4, 0, 0] \text{ m/s}$ | $\frac{\partial p}{\partial n} = 0$ |
| Outlet | Outlet | $\frac{\partial \mathbf{u}}{\partial n} = 0$ | Advective with $L_{\text{inf}} = 0.5 \text{ m}$ |
| Side walls | Wall | $\mathbf{u} = 0$ | Slip condition |
| Fan casing | Wall | $\mathbf{u} = 0$ | $\frac{\partial p}{\partial n} = 0$ |
| Fan blades | Moving wall | $\mathbf{u} = \omega \times \mathbf{r}$ | Fixed flux |
| AMI interfaces | Cyclic AMI | Matched values | Matched values |

To model the rotating auxiliary fan, the Arbitrary Mesh Interface (AMI) method was employed with a rotating motion defined by:

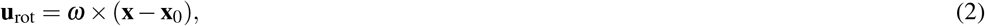

where ω = [0, −360, 0] rad/s is the angular velocity vector and **x**_0_ is the rotation origin. The angular velocity was chosen based on the specifications of the physical fan used in our experiments (Delta Electronics, 2024). The fan rotation was implemented using the dynamic mesh capabilities of OpenFOAM with solid body motion solver.

The turbulence was modeled using the *k*-ω SST formulation with stabilization parameters α_*k*1_ = 0.85 and α_*k*2_ = 1.0. Turbulent viscosity was limited with *k*_max_ = 10 m^2^*/*s^2^ and ω_max_ = 1×10^12^ s^−1^ to ensure numerical stability. The PIMPLE algorithm was employed for pressure-velocity coupling with 3 outer correctors and under-relaxation factors of 0.3 for pressure and 0.7 for velocity. The simulation was initialized with quiescent conditions (**u** = 0 throughout the domain) and run for 20 seconds of physical time at a rate of 100 Hz. The initial transient period of approximately 5 seconds, corresponding to the flow development from quiescent conditions, was excluded from subsequent analysis.

### Experimental Flies

All experiments used female progeny from a cross between male wild-type Heisenberg-CantonS flies and virgin female Norpa;Orco-Gal4;UAS-CsChrimson flies, as detailed in (Stupski and van Breugel, 2024). Female flies were selected for their larger body size, which improves tracking robustness, and their reduced susceptibility to desiccation in the wind tunnel environment.

Flies were maintained following established protocols (Stupski and van Breugel, 2024) on a 16:8 light:dark cycle at 25°C and 60% relative humidity. They were reared on in-house fly media based on the traditional Caltech recipe (cornmeal, sucrose, dextrose, yeast, and 2-acid medium (Lewis, 1960)), with supplemental yeast added to each bottle. Each bottle was also supplemented with 400 µL of 40 mM all-trans-Retinal (ATR; R2500, Sigma Aldrich) dissolved in sucrose solution. Following eclosion, flies were transferred to fresh ATR-supplemented media for at least 48 hours prior to experimentation. Each experimental night, 15 F1 Orco*>*CsChrimson females (2–5 days old) were placed in empty test tubes with moist Kimwipes to prevent desiccation. Flies were collected in the morning, starved for 6–8 hours, then introduced into the wind tunnel in the evening for overnight free-flight recording.

#### Trajectory Inclusion Criteria and Data Filtering

Trajectory analysis included only flights with at least one second of tracking after the trigger event. We excluded trajectories containing any ground speed measurements exceeding 2.0 m/s or positional data points outside the physical tunnel bounds, which are often associated with flies walking along walls. To further filter out walking behavior, we discarded trajectories where total displacement was less than 0.02 m or mean ground speed fell below 0.15 m/s. Because the tracking system could not reliably maintain individual fly identity across the overnight recording session, each trajectory was treated as statistically independent. Data collection spanned 43 nights (approximately 645 flies).

## RESULTS

### Auxiliary fans within wind tunnel induce distinctive flow regions

To investigate the role of wind history during in-flight olfactory search, we added two auxiliary fans downwind within our wind tunnel (Figure 1A-C), with one on each side wall. The auxiliary fans were programmed to alternatively cycle on and off at 20-minute intervals, with an additional 20-minute period where both fans were off. We refer to these auxiliary fans as “gusts” to distinguish them from the mean bulk flow generated by the upwind multi-fan array. Upon analysis, flight responses to either auxiliary fan were found to be largely symmetric; therefore, we mirrored the data to increase the interpretability of results (see Supplemental Figure S1 for details). Throughout the experiment, flies entered a predetermined trigger volume (Figure 1B) and received either a 0.675s red light stimulus (“flash”) or no stimulus (“sham”). Prior work (in the same wind tunnel) has shown that this protocol replicates behavior similar to that observed in plume tracking experiments with real odor, with minimal behavioral artifacts (Stupski and van Breugel, 2024).

We simulated our wind tunnel using computational fluid dynamics (CFD) to visualize the resulting flow dynamics (Figure 1C). A 3-D streamline reconstruction of the flow field at simulation time *t* = 10s demonstrates smooth laminar flow in the upwind region and a more dynamic, unsteady region downstream arising from flow interaction with the auxiliary fans (Figure 1D). The (*x, y*) velocity field (Figure 1E) further illustrates the strong crosswind component introduced by an active auxiliary fan. To validate the model, wind measurements were collected at 6 different *x*-positions along the *y*-centerline for approximately 2 minutes per location, and the mean measurements were compared to mean CFD wind speeds sampled over a corresponding volumetric region for 5 ≤ *t* ≤ 20s (Figure 1F). Due to the mesh size, we were able to take a larger sample of virtual measurements within our volumetric region, as can be seen in Figure 1F. General agreement was observed between physical and simulated results. We note some variation, likely attributable to flow interactions introduced by the physical measurement apparatus that were absent in the simulation.

### Instantaneous wind experience predominantly drives surge behavior during olfactory stimulus

From our physical measurements and CFD flow reconstruction (Figure 1D-F), it was clear that flies could encounter very different wind conditions depending on their position within the tunnel. To better understand the behavioral influence of wind experience during an olfactory stimulus, we segmented flies into three groupings (“upwind”, “gust zone”, and “downwind”) based on their mean *x*-position during the olfactory stimulus (0 ≤ *t* ≤ 0.675s; Figure 2A-C). To estimate the wind each fly actually experienced, we initialized each trajectory within the 3-D CFD instantaneous velocity field at five different simulation start times (5 ≤ *t* ≤ 20s) and averaged the reconstructed wind velocities. The resulting *x*- and *y*-velocity kernel density estimates (KDEs) for the first half of the red light stimulus (0 ≤ *t* ≤ 0.3s) are shown in Figure 2B. Flies in the upwind region (orange) primarily experienced the mean bulk flow, with *x*-velocity ≈ −0.4m/s and minimal crosswind component. Flies in the gust zone (cyan) encountered a much broader range of wind velocities, with crosswind gusts approaching ≈1.25m/s. Flies classified as downwind (purple) also experienced variable wind, but without the strong lateral gust. Representative example trajectories illustrate the distinct behavioral responses associated with each grouping (Figure 2D).

**Figure 2:**
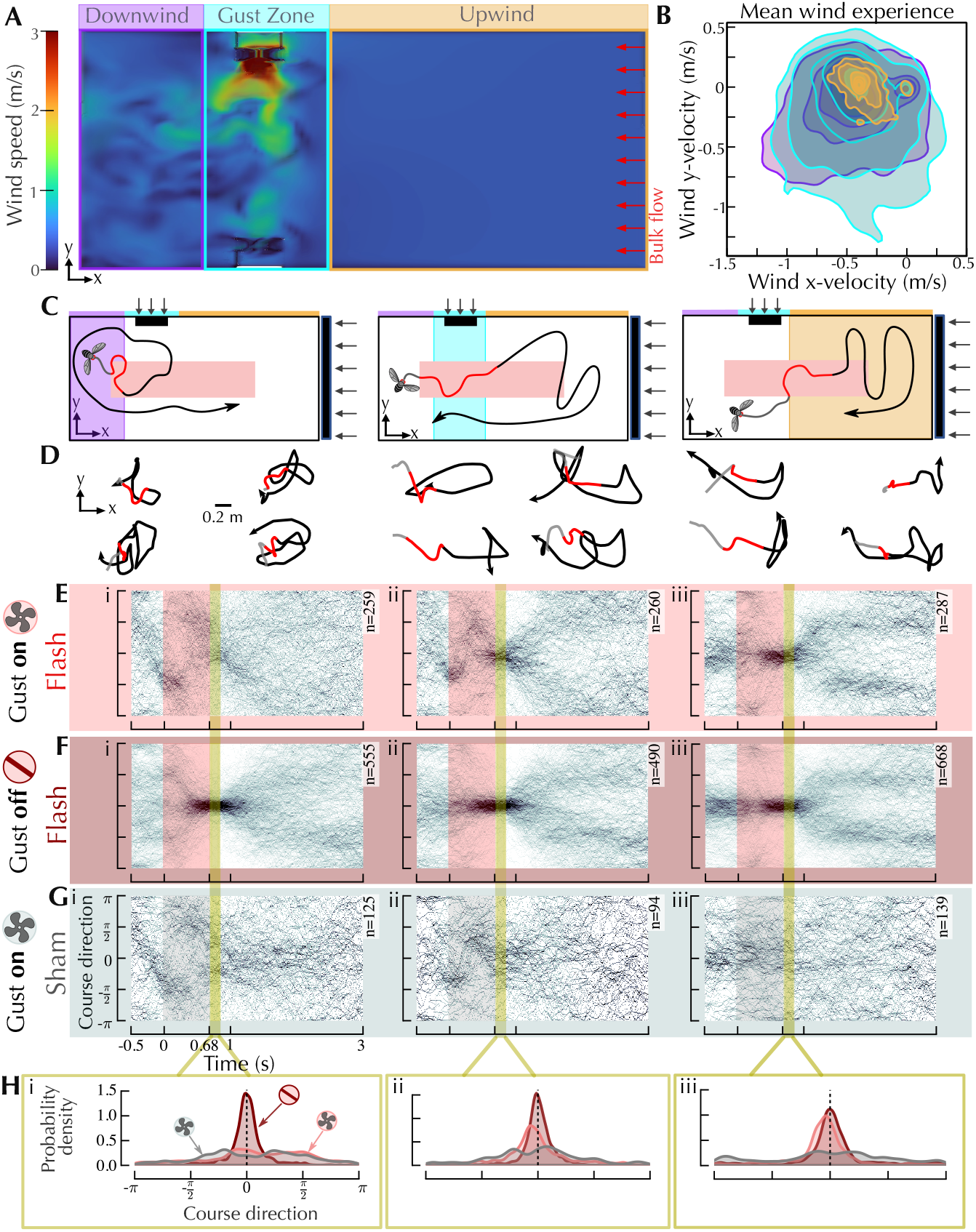
Olfactory surge direction is largely governed by instantaneous wind experience. (A) 2-D (*x, y*) slice of the instantaneous wind speed profile (as in Figure 1E) with fly groupings indicated. Flies were assigned to “upwind”, “gust zone”, or “downwind” categories based on mean *x*-position during the olfactory stimulus (0 ≤ *t* ≤ 0.675s). (B) 2-D kernel density estimates (KDEs) of the wind velocity experienced by flies in each grouping during the first half of the red light stimulus (0 ≤ *t* ≤ 0.3*s*). For each fly, the experienced wind velocity was sampled from the instantaneous 3-D CFD velocity field based on the fly’s position, averaged across 5 different CFD start times to account for gust phase variability. Each set of contours (purple, green, orange) represents the estimated joint probability density of mean x-velocity and mean y-velocity for the upwind, gust zone, and downwind groupings described in (A). Contours are drawn at 5 equally spaced density levels, with the outermost contour corresponding to 5% of the peak density; inner contours indicate regions of progressively higher data concentration. Note that axes represent wind velocity components, not spatial position. (C) Schematic highlighting the grouping criteria; colored regions correspond to upwind, gust zone, and downwind areas. The red zone indicates the approximate trigger region for the red light stimulus. (D) Example trajectories for flies assigned to each grouping. (E-G) Aggregate course direction for (E) flies that received an olfactory stimulus while the gust was on, (F) flies that received an olfactory stimulus while the gust was off, and (G) sham events while the gust was on. A course direction of 0 radians indicates upwind flight while *π*/2 radians indicates flight toward the auxiliary fan. (H) Probability density estimates of course direction immediately after olfactory stimulus offset (0.675 ≤ *t* ≤ 0.8s) for scenarios i-iii.

To better visualize differences in behavior as a function of flow condition, we compared the aggregate course direction for flash and sham events across the three groupings while an auxiliary gust was present (Figure 2E), while the fans were off (Figure 2F), and for shams while a gust was on (Figure 2G). Flies in the downwind region during olfactory stimulus exhibited a typical upwind surge when the auxiliary fans were off (Figure 2Fi), whereas downwind flies exposed to an active gust showed a sustained change in course direction before and during the red light stimulus, with no clean upwind surge. Importantly, this course-direction perturbation was observed in both flash and sham flies (Figure 2E-F), indicating that the maneuvers occurring approximately −0.5 ≤ *t* ≤ 0.5s are largely wind mediated rather than olfactory driven.

For upwind flies, surge behavior was consistent regardless of whether the gust was on or off (Figure 2Eiii, Fiii), confirming that flies in the upwind laminar region were not strongly influenced by the downwind gust perturbation. Flies in the gust zone, however, exhibited a distinctive three-phase course correction during olfactory stimulus: an initial crosswind deflection caused by the gust, followed by a compensatory turn back toward the gust direction, and finally an upwind surge (Figure 2Eii). The probability density of course direction angle immediately after olfactory stimulus offset (Figure 2Hi-iii) further shows that both upwind and gust zone flies exhibit an upwind surge with a slight directional bias away from the gust of approx. -9 degrees (Figure 2Hii-iii). This upwind bias is absent in the corresponding sham conditions, indicating that the olfactory driven upwind surge response is robust to being temporarily blown off course.

### Flies preferentially return to the flow region experienced during olfactory stimulus

To explore how wind experience during an olfactory stimulus influences subsequent search behavior, we examined the positional and course direction preferences of each grouping during the search phase after surge behaviors had largely ended (1.3 ≤ *t* ≤ 3s; Figure 3A). Occupancy maps (Figure 3B) show the (*x, y*) positions of flies during the search phase, revealing distinct spatial preferences across the upwind, gust zone, and downwind groupings. Flies that received an olfactory stimulus while the gust was off (Figure 3Bii) largely displayed the expected upwind positional preference, whereas flies that received an olfactory stimulus while the gust was on showed greater occupancy in the regions near where they originally received the olfactory stimulus (Figure 3Bi). While spatial preferences were most pronounced for flies that received an olfactory stimulus while the gust was active, a weaker trend was also evident for sham flies and for flies that received an olfactory stimulus while the gust was off, indicating a partial contribution from both baseline positional biases and olfactory drive.

**Figure 3:**
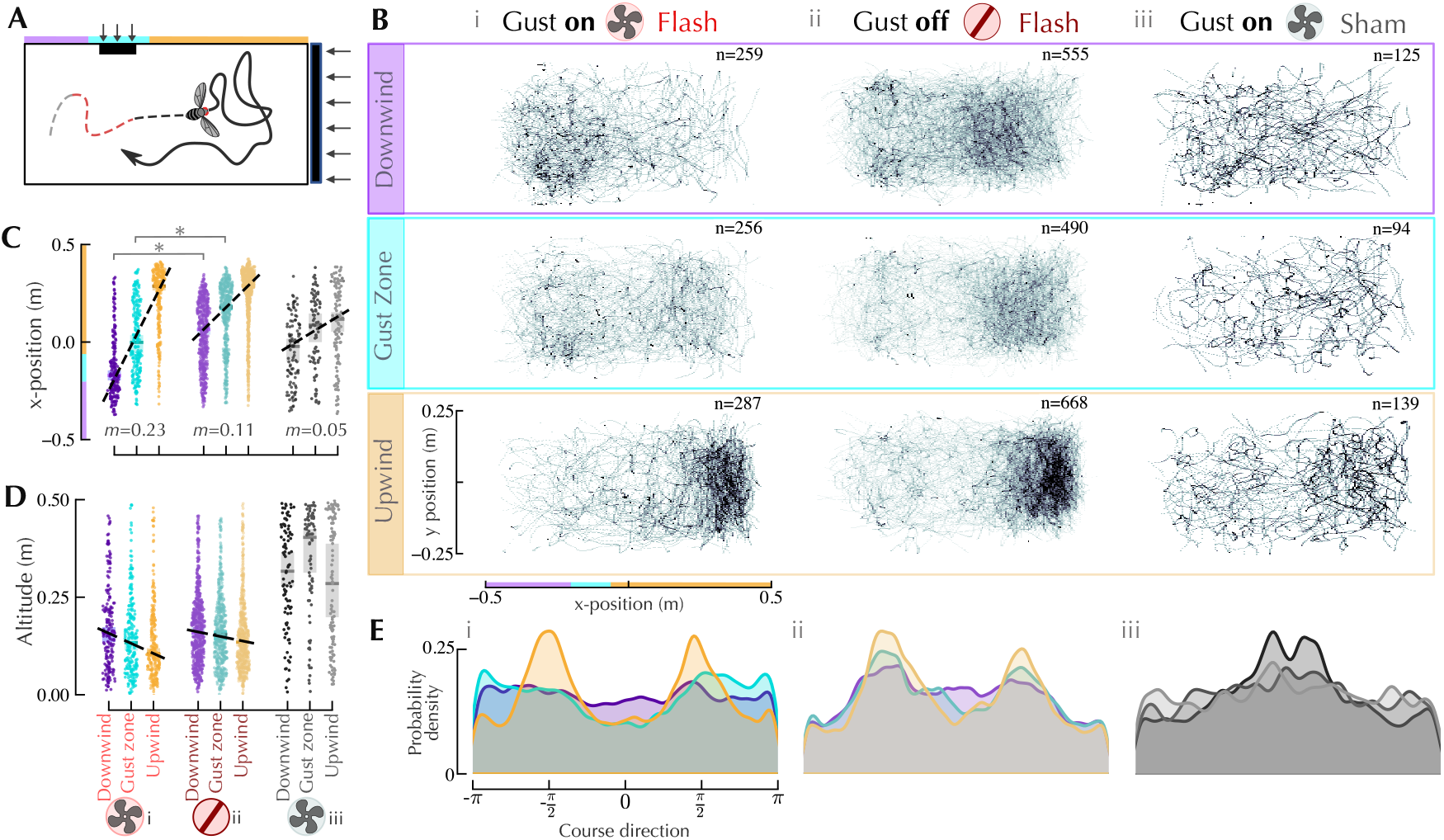
Wind experience during olfactory stimulus shapes where insects subsequently choose to search. (A) Schematic demonstrating trajectory search phase. As in Fig 2, trajectories were grouped as “upwind”, “gust zone”, or “downwind” based on mean *x*-position during olfactory stimulus (0 ≤ *t* ≤ 0.675s). The trajectory shown exemplifies a “gust zone” fly. (B) Occupancy maps showing (*x, y*) positions of flies during the search phase (1.3 ≤ *t* ≤ 3s). Total trajectory counts for each grouping are shown in the upper right-hand corner. (C) Mean *x*-positions for trajectories across the upwind, gust zone, and downwind groupings based on whether flies: (i) received an olfactory stimulus while the gust was on, (ii) received an olfactory stimulus while the gust was off, or (iii) were shams while the gust was on. Black dashed line corresponds to the median across all trajectories within each grouping; values below indicate the line slope. Asterisks above indicate significant differences at the *p <* 0.001 level (Mann-Whitney U test). (D) Mean altitudes for the same groupings as in (C). (E) Probability density estimates of course direction across groupings for 1.3 ≤ *t* ≤ 3s.

To quantify these trends, we computed the mean *x*-position for each trajectory across the three groupings under three scenarios: (i) flash while gust on, (ii) flash while gust off, and (iii) sham while gust on (Figure 3C). We then computed the median slope for each grouping’s aggregate x-positions based on whether they were downwind, upwind, or in the gust zone. In sham flies, the small positive slope of *m* = 0.05m across groupings indicates that flies categorized as upwind were more likely to be upwind during their search phase than flies in the gust zone or downwind groups (Figure 3Ciii). The median x-position slope roughly doubled (*m* = 0.11m) for flies that received an olfactory stimulus with the gust off (Figure 3Cii), indicating that flies grouped as upwind when receiving their olfactory stimulus remained further upwind than their gust zone and downwind counterparts. When the gust was active and flies received a red light stimulus, the trend was further amplified (*m* = 0.23m, Figure 3Ci), with a much larger spread in mean x-position as a function of where the flies initially received their olfactory stimulus. In this scenario, a large proportion of flies that received their olfactory stimulus while downwind chose to stay downwind, whereas flies that received their olfactory stimulus near the gust zone also searched closer to the gust zone on average. Downwind and gust zone flies that received an olfactory stimulus showed significantly different x-positional preferences when the auxiliary fan was either on or off, with both groupings remaining mostly upwind if the gust was off versus staying more downwind if the gust was active (Mann-Whitney U test, p!0.001, Figure 3C). Taken together, these results indicate that, after accounting for baseline positional artifacts, flies that experienced a more complex windscape during their olfactory stimulus preferentially chose to remain in, or return to, the flow region where the olfactory stimulus occurred.

We also examined altitude and course direction during the search phase. A small altitude trend was observed across upwind, gust zone, and downwind flies that received an olfactory stimulus, where flies were more likely to exhibit lower altitudes if they were closer toward the upwind section of the tunnel (Figure 3Di-ii). This altitude effect is consistent with prior reports of odor triggered altitude reduction (Stupski and van Breugel, 2024; Van Breugel and Dickinson, 2014). In comparison, no clear reduction in altitude was present in sham flies. Course direction during the search phase varied systematically: upwind flies showed clear crosswind casting (Figure 3Ei-ii), gust zone flies showed a downwind casting bias (Figure 3Eii), whereas downwind flies showed no directional preference. We note that the positional behavior of flies in the downwind grouping was less consistent even when the gust was off (Figure 3Biii), suggesting residual positional artifacts. As such, we focus our subsequent analyses on the upwind and gust zone groups.

### Gust-displaced flies are more likely to explore the gust zone during subsequent search

Our analysis of flies that received an olfactory stimulus during an active gust revealed a bimodal search response. Flies either chose to stay upwind and perform a stereotypical casting maneuvers, or they chose to turn downwind back toward the gust. To further characterize the behavior of these flies, we separated gust zone flies that remained upwind (*x* ≥ 0.1m) from those that traveled downwind after olfactory stimulus offset (Figure 4A-D). Both groups initially traveled out of the gust zone following olfactory stimulus offset, with a divergence occurring around *t* = 1.2*s*, when gust-returning flies made an active course reversal back toward the downwind region (Figure 4C). Flies that returned toward the gust did not exhibit a clear casting pattern (Figure 4B,D), which may reflect physical wind tunnel boundary constraints limiting the full expression of this behavior, or may be a consequence of the strong, directionally variable wind they were experiencing. The course direction of flies that remained upwind versus returning downwind shows a clear crosswind versus downwind preference, respectfully (Figure 4D).

**Figure 4:**
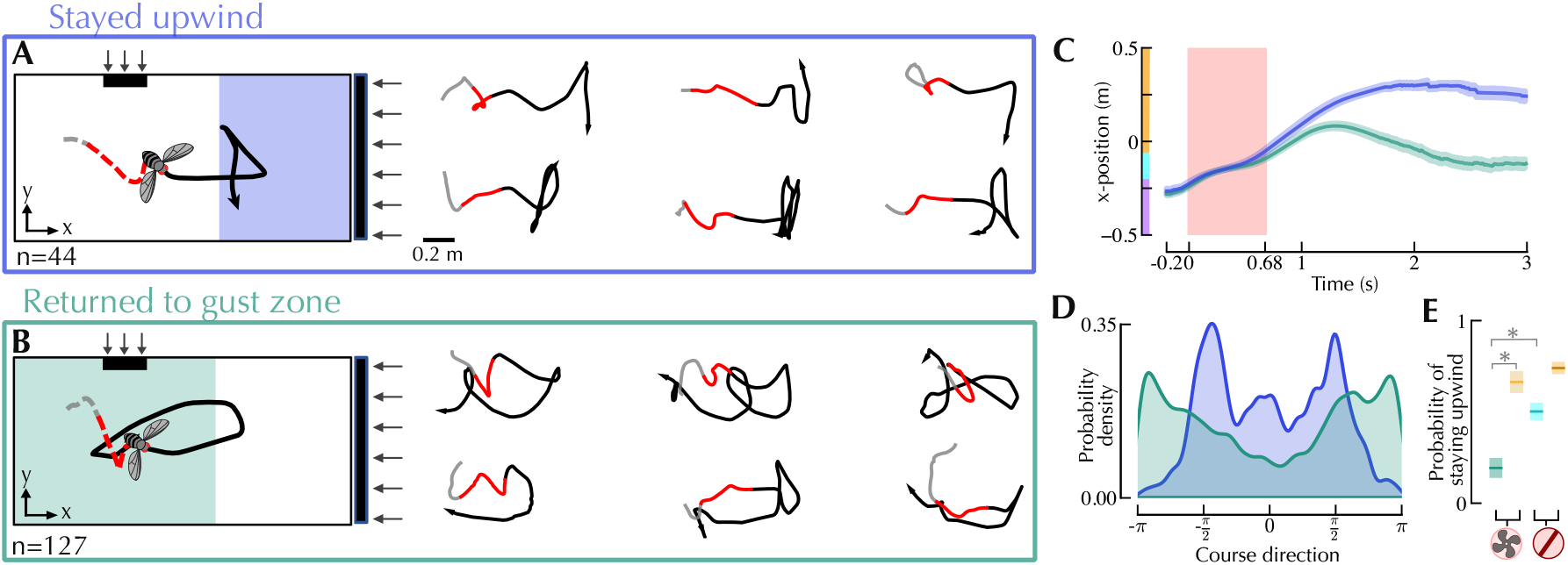
Flies displaced by a gust during olfactory stimulus are likely to explore the gust zone during search. (A) Schematic and example trajectories for gust zone flies that remained upwind (*x* ≥0.1m) after the olfactory stimulus. Flies with an initial downwind course before olfactory stimulus onset, and trajectories with less than 0.2s of pre-flash observation were excluded. (B) Schematic and example trajectories for flies that traveled downwind during the search phase. (C) Mean *x*-position over time for flies that remained upwind versus those that showed a downwind preference. Shading represents 95% confidence intervals. (D) Course direction probability density for trajectories in (A-B) during 1.3 ≤ *t* ≤ 3s. (E) Probability that a fly remained upwind (*x >* 0.1*m*), as a function of its flow experience and mean *x*-position during the olfactory stimulus. Comparative groupings are shown for flies that were either in the gust zone (teal) or upwind (gold) when they received their olfactory stimulus and the auxiliary fan was active, as well as for flies that were in the gust zone (cyan) or upwind (brown) but received an olfactory stimulus while the gust was off. Total populations from left to right: *n* = 197, *n* = 235, *n* = 426, *n* = 616. Asterisks indicate significant differences at the *p <* 0.001 level (proportion *z*-test). Shading throughout indicates 95% confidence intervals.

To examine how frequently flies in the gust zone were likely to re-explore the gust area, we estimated the probability of a fly choosing to remain upwind and quantified uncertainty via Bernoulli confidence intervals, with proportion *z*-tests used to assess significance. This probability was estimated across four groups: upwind and gust zone flies that received an olfactory stimulus while the gust was active, as well as upwind and gust zone flies that received an olfactory stimulus while the auxiliary fan was inactive (Figure 4E). The probability of remaining upwind was substantially lower for flies that experienced a gust during the olfactory stimulus when compared to flies that were upwind of the gust. Furthermore, gust zone flies whose olfactory stimulus occurred while the fans were off were significantly more likely to remain upwind than flies that received both olfactory and gust stimuli. Examination of mean trajectory positions, velocities, and wind experience in the period immediately prior to, and during, the red light stimulus (−0.2 ≤ *t* ≤ 0.675*s*) revealed some slight differences, but the small sample size and lack of heading data make it difficult to determine a causal effect of search preference between flies that chose to stay upwind versus those that returned to the gust zone (Supplemental Figure S2). Regardless, these population level findings provide evidence that flies keep track of the wind experienced during an olfactory stimulus and use this information to guide spatial exploration during the subsequent search phase.

## DISCUSSION

Our results demonstrate that wind experience during an olfactory stimulus influences both the immediate surge response and subsequent spatial search in freely flying *Drosophila melanogaster*. Flies displaced by a crosswind gust at the moment of odor contact executed a three-part course correction: crosswind deflection, compensatory turn, and upwind surge. Despite potential displacement by the gust, these flies still largely turned to surge upwind after leaving the gust zone, indicating that the surge response is resilient to transient flow perturbations and more likely mediated by instantaneous wind rather than wind history. Notably, these same flies were significantly more likely to return to the gust zone during their post-olfactory stimulus search phase, even though the wind conditions may have been physically challenging to fly in. These positional preferences cannot be fully explained by artifacts of our *x*-position-based grouping criterion, nor by pre-olfactory stimulus behavioral differences. Together, our results suggest that wind is estimated and stored as a distinct navigational state, in addition olfactory experience. Furthermore, we provide evidence that wind memory actively shapes where flies choose to search after receiving an olfactory stimulus, while course direction during surge and search may be more reflexive to instantaneous flow.

Prior work has established that insects retain short-term memory for odor encounter history in flying *Drosophila* and *Aedes aegypti* (Pang et al., 2018b), while encounter frequency modulates locomotor gating in walking flies (Álvarez Salvado et al., 2018). Our results extend this picture by showing that wind is a state tracked in addition to odor experiences. The fact that gust-exposed flies chose to revisit the gust zone even in the absence of an ongoing odor signal suggests they are using retained wind information to direct spatial exploration. This is consistent with the broader view, supported by computational modeling and prior empirical evidence (Singh et al., 2023; Grü nbaum and Willis, 2015; Pang et al., 2018a), that multi-timescale sensory integration, including memory for both odor and flow conditions, is a core feature of robust plume-tracking.

An open question raised by our results is *how* flies implement the decision to return to the gust zone versus remaining upwind. Our current analysis tracks positional and course direction preferences at the population level, but the key decision likely plays out at the level of individual biomechanical maneuvers. Specifically, whether and when a fly makes a directed saccadic turn toward the gust zone after its upwind surge terminates. Future experiments incorporating heading angle as a primary readout, rather than course direction alone, would allow us to distinguish between active reorientation toward the gust and passive drift caused by strong flow perturbations. Combining such heading-resolved tracking with optogenetic silencing of specific olfactory or mechanosensory pathways could further establish which sensory modalities and brain circuits are required to encode and recall wind history information. Furthermore, the results of this study raise the question of how flies know where to go after receiving their stimulus. Future studies would be needed to tease out the extent to which flies are using visual landmarks or odometry for guidance versus solely following a wind-gated reflex to return to the area they received their olfactory stimulus. More broadly, our results motivate further exploration of the neural architecture underlying sensory processing of ambient wind in olfactory search.

## AUTHOR CONTRIBUTIONS

J.H. and F.v.B. conceived the presented idea; J.H. collected and analyzed the data; J.H. created the CFD simulation; A.P.L. collected the physical wind measurements; J.H. and F.v.B. wrote the main manuscript.

## DATA AVAILABILITY

Data will be made available upon publication.

## ACKNOWLEDGMENTS

We are grateful for feedback from S. David Stupski during experimental design and for assistance with data collection for this project. We also thank Aditya Nair for his suggestions to improve the CFD modeling.

