## Supplemental Material for "Wind history shapes olfactory search response in free flying *Drosophila melanogaster*"

### SUPPLEMENTARY MATERIAL

#### Mirroring Left and Right Gust

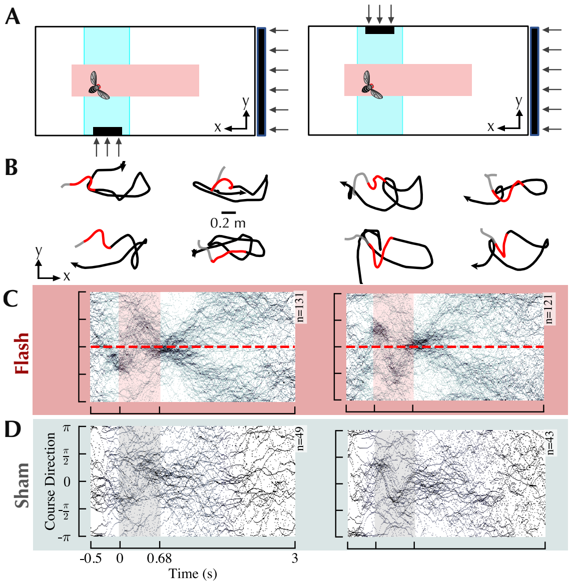


Supplementary Figure S1: **Aggregate course directions were symmetric for the left and right gust scenarios.**

(A) Schematic depicting the subset of flies chosen based on their mean position during the red light stimulus. In either scenario, either the auxiliary fan at $y = -0.25m$ or at $y = 0.25m$ was turned on. (B) Example (x,y) trajectories for the two flow scenarios. (C) Aggregate course direction plots for flies that received a red light stimulus in the two gust scenarios. (D) Same as in (C) but for shams.

### Separating gust zone flies that stayed upwind or went downwind show no clear preemptive behavioral differences

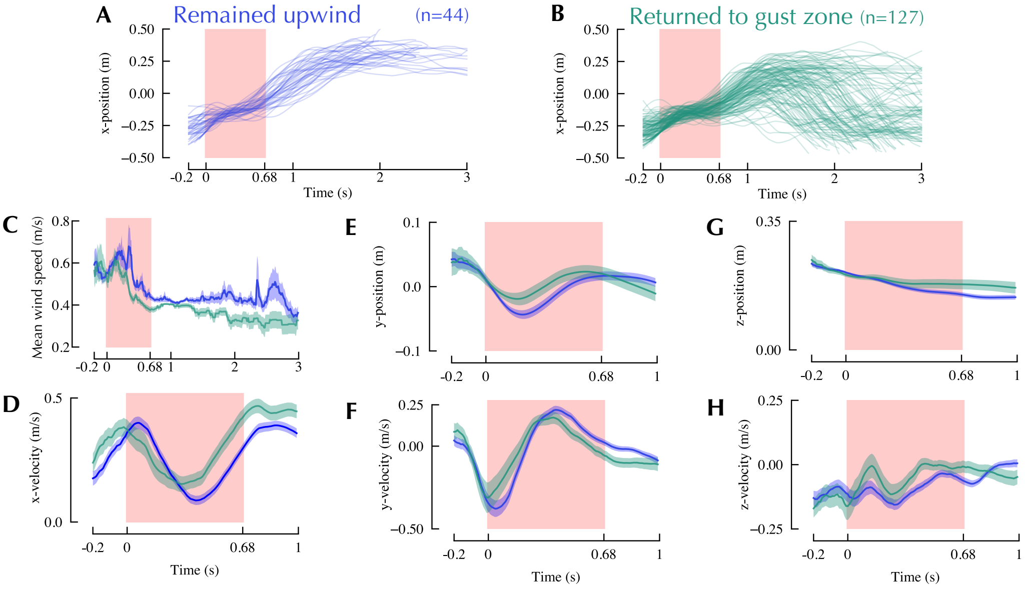


Supplementary Figure S2: **Fly velocities and mean reconstructed wind speed showed no clear causes for up- versus downwind search response.**

(A) X-position over time for all flies that maintained an x-position ¿ 0.1m during the search phase of their trajectory. (B) X-position over time for flies that did not meet the criteria in (A). (C) Mean reconstructed wind experience (found using CFD wind velocity field) for gust zone flies that either stayed upwind during search (n=44) or went downwind (n=127), as depicted in Figure 4. (D) Mean x-velocity for the two groups. (E) Mean y-position for the two groups. (F) Mean y-velocity for the two groups. (G) Mean z-position for the two groups. (H) Mean z-velocity for the two groups.
